# *Galleria mellonella* as an Insect Model for *P. destructans*, the Cause of White-Nose Syndrome in Bats

**DOI:** 10.1101/244921

**Authors:** Chapman Beekman, Lauren Meckler, Eleanor Kim, Richard J. Bennett

## Abstract

*Pseudogymnoascus destructans* is the fungal pathogen responsible for White-nose Syndrome (WNS), a disease that has killed millions of bats in North America over the last decade. A major obstacle to research on *P. destructans* has been the lack of a tractable infection model for monitoring virulence. Here, we establish a high-throughput model of infection using larvae of *Galleria mellonella*, an invertebrate used to study host-pathogen interactions for a wide range of microbial species. We demonstrate that *P. destructans* can kill *G. mellonella* larvae in an inoculum-dependent manner when infected larvae are housed at 13°C or 18°C. Larval killing is an active process, as heat-killed *P. destructans* spores caused significantly decreased levels of larval death compared to live spores. We also show that fungal spores that were germinated prior to inoculation were able to kill larvae 3–4 times faster than non-germinated spores. Lastly, we identified chemical inhibitors of *P. destructans* and used *G. mellonella* to evaluate these inhibitors for their ability to reduce virulence. We demonstrate that two chemicals, trifluoperazine and amphotericin B, can effectively block larval killing by *P. destructans* and thereby establish that this infection model can be used to screen biocontrol agents against this fungal pathogen.

## Introduction

*Pseudogymnoascus destructans* is the fungal species responsible for White-nose Syndrome (WNS), a disease currently devastating bat populations across North America. *P. destructans* is a psychrophilic fungus and colonizes susceptible bat species during hibernation, causing depletion of energy stores and death of the host. Since it was first discovered in New York State in 2006, WNS has spread to 31 US states and 5 Canadian provinces [1]. This rapid spread, combined with a mortality rate approaching 100% for several species, has led to an estimated 6 million bats being killed by WNS [1]. As a result, one of the most common bat species in the North-East US, the little brown bat (*Myotis lucifugus*), is now threatened with regional extinction. The loss of bats can harm both local ecosystems and agriculture as they play a crucial role in controlling insect pest populations, and it has been estimated that bat populations lost to WNS could cost the agricultural industry as much as $23 billion per year [2].

Despite the impact of WNS, a clear understanding of the factors that allow *P. destructans* to infect its host remain elusive. Studies have suggested several attributes may be important for fungal virulence including the production of small molecule effectors [3], protease secretion [4, 5], lipid utilization [6], as well as the fungal heat shock response, cell wall remodeling, and micronutrient acquisition [7]. A significant obstacle to evaluation of these hypotheses has been the lack of a tractable infection model. The psychrophilic nature of *P. destructans* (maximum growth temperature of ~18 °C) has made standard mammalian infection models unfeasible. Laboratory-based WNS models using live bats have been utilized [8, 9], but require specialized equipment and long infection timelines, and also are impractical for high-throughput studies.

The lack of an accessible infection model for *P. destructans* has also limited the testing of therapeutic agents to treat WNS. Studies have identified several agents that can inhibit *P. destructans* growth on laboratory media [10–12], yet it is difficult to test these in a natural setting. Treating a live infection entails additional complications that are not present during growth on laboratory media, such as host drug toxicity or drug degradation by the host. Additionally, available carbon sources and growth conditions can affect fungal drug susceptibility [13, 14]. Treatment of WNS also poses unique challenges given that infection occurs only in hibernating bat populations, often located in remote habitats. Development of a simple host model of *P. destructans* infection would therefore be of considerable value and could accelerate the identification of an effective treatment for WNS.

For several fungal pathogens, larvae of the greater wax moth, *Galleria mellonella*, have provided a simple yet effective alternative to mammalian infection models. *G. mellonella* has been employed to study virulence in human fungal pathogens including *Candida* [15–17], *Aspergillus* [18, 19] and *Fusarium* [20, 21] species. Importantly, many results obtained using *G. mellonella* reproduce findings from mammalian infection studies [15, 19, 21, 22], indicating that larval infection shows parallels with that in higher eukaryotes. Though insects lack an adaptive immune system, their immune response closely resembles the mammalian innate immune response at both a structural and functional level [23]. *G. mellonella* hemocytes are functionally analogous to mammalian phagocytes and generate reactive oxygen species (ROS) for microbial killing [24, 25]. These features make *G. mellonella* a relevant model for WNS, as evidence suggests hibernating bats can mount an innate immune response to *P. destructans* but are unable to activate adaptive immunity [26–28]. To our knowledge, no previous studies using *G. mellonella* have been conducted at temperatures below 20**°**C, yet larvae can be maintained at such temperatures making their use as a *P. destructans* infection model an attractive possibility.

Here, we examine the feasibility of using *G. mellonella* larvae as an invertebrate model for infection with *P. destructans*. Our results indicate that live *P. destructans* spores, but not heat-killed spores, are lethal to *G. mellonella* larvae, and that killing is augmented if spores are induced to germinate prior to inoculation. We also perform a screen to identify novel chemical inhibitors of *P. destructans* growth, and evaluate their efficacy during infection. These experiments establish that insect larvae can be used as a high-throughput model for *P. destructans* enabling the screening of potential treatments for WNS.

## Materials and Methods

### Strains and culture conditions

*Pseudogymnoascus destructans* strain 20631–21 (ATCC stock: MYA-4855) was used for all experiments. *P. destructans* cultures were cultured on yeast extract-peptone-dextrose (YPD) medium at 13**°**C prior to spore collection.

### *G. mellonella* virulence assays

*G. mellonella* larvae were obtained from Vanderhorst Wholesale (St. Marys, OH). Prior to infection, larvae were stored at 13**°**C and used within 1 week of delivery. For all experiments, inoculums were prepared by harvesting *P. destructans* spores from 2–3 week-old cultures grown on YPD plates using a solution of 0.05% Tween-20 (Sigma) and rubbing the surface of the plates with a glass spreader to release spores. Spores were isolated from larger hyphae by filtering through a layer of sterile miracloth, centrifuged and resuspended in 5 mL sterile phosphate-buffered saline, pH 7.4 (PBS). Spores were counted on a light microscope (Leica DM750) using a hemocytometer and adjusted to the desired concentration. Heat-killed spores were prepared by incubating inoculums for at least 30 min at 65**°**C and re-cooling briefly on ice prior to injection. For pre-germination experiments, *P. destructans* spores were adjusted to 1×10^7^ cells/mL in liquid YPD and incubated at 13**°**C in a shaking incubator (200 rpm) for the specified time period. Germination of spores was assessed by placing 10 µL of the inoculum on a glass slide or hemocytometer and counting the number of germ tubes versus un-germinated spores. Approximately 400 spores were counted in two independent experiments. Inoculums were then prepared by adjusting spores to 1×10^8^ cells/mL in sterile PBS. The concentration of spores in each inoculum was confirmed by re-counting spores prior to injection. Worms were randomly selected for each experiment and those showing signs of discoloration were discarded. Each worm was injected with 10 µL of inoculating solution just below the second to last left proleg using a 10 µL glass syringe with a 26S gauge needle (Hamilton, 80300). Infected larvae and controls were maintained at 13**°**C and checked daily. Worms were recorded as dead if no movement was observed upon contact. For antifungal experiments, *P. destructans* spores were pre-incubated in YPD for 6 h then resuspended in PBS containing the antifungal compound being tested. Spores were re-counted prior to injection and adjusted to 1×10^8^ cells/mL if necessary. For antifungal experiments, spores were exposed to each compound for up to 2 hours prior to injection.

### Phenotype Microarray (PM) screen

*P. destructans* spores were harvested, washed in PBS, counted, and adjusted to 1×10^6^ cells/mL in liquid YPD. Next, 100 µL of the spore solution was used to inoculate each well of PM plates 21–25 (Biolog, Hayward, CA). Each well contains one of 120 different chemical compounds at 4 different concentrations [29]. Plates were incubated at 13**°**C and growth in each well was monitored daily by measuring optical density (OD595 nm) in a Synergy HT plate reader (Biotek, Winooski, VT) for 11 days. OD readings were compiled to construct growth curves which were analyzed using DuctApe™ software [30] to convert each growth curve into an “activity index” (0–9) representing relative growth of *P. destructans* in response to each chemical. Data shown represents the average activity indices from 2 independent experiments.

### Comparison of *P. destructans* and *C. albicans* PM data

Data from the PM drug screen on *P. destructans* was compared with published data obtained for *C. albicans* using the same PM drug panel and DuctApe analysis software [31]. Inhibitory compounds were defined as those producing an activity index of 0 at any concentration. Individual compounds were compared using the BioVenn [32] webtool (http://www.biovenn.nl/index.php, ©2007 - 2017 Tim Hulsen). Compounds were categorized by “mode of action” and each category was evaluated for enrichment of inhibitory compounds (Fishers exact test, Graphpad Prism v.5).

### Determination of minimum inhibitory concentrations (MICs) of select compounds

*P. destructans* spores were harvested and adjusted to 1×10^6^ spores/mL in YPD solutions containing test compounds. These solutions were then used to inoculate individual wells of clear plastic 96-well tissue culture plates (100 µL/well, 6 replicate wells/condition). Plates were incubated at 13**°**C and growth (OD595) measured after 240 h using a Synergy HT plate reader (Biotek). OD values from each of the 6 replicate wells were averaged and then divided by the average growth in control wells containing spores in YPD to determine the fraction of growth. The MIC was defined as the lowest concentration of each compound able to reduce *P. destructans* growth by at least 80%.

### Evaluation of compounds for fungicidal activity

Harvested *P. destructans* spores were adjusted to 1×10^6^ spores/mL in YPD containing the test compound and incubated at 13**°**C for 24 h. Next, serial dilutions of the spore solutions were made in YPD and 2 µL of each dilution spotted onto YPD plates, incubated at 13**°**C for 1 week and imaged using a Chemidoc imaging system (BioRad).

## Results

### Evaluation of *G. mellonella* as a suitable host for *P. destructans*

To test if *P. destructans* can establish an infection in *G. mellonella*, larvae were injected with 10^4^, 10^5^, or 10^6^ fungal spores or an equal volume of phosphate-buffered saline (PBS). Injected larvae were incubated at either 13°C or 18°C, as appropriate temperatures for growth of *P. destructans*. At 18°C, the upper limit for growth of *P. destructans*, increased inoculum size correlated with increased killing of *Galleria* larvae (Fig. 1A). In contrast, at 13°C, only the highest inoculum of 10^6^ spores/larva demonstrated increased lethality above the PBS-injected control group (p = 0.0167, log-rank test, Fig. 1B). At the highest inoculum, the rate of killing was similar at both 13°C and 18°C, with 50% mortality reached ~20–25 days after infection. Larvae injected with the highest fungal inoculum also showed an increase in pigmentation within 2 weeks of injection (Fig. 1C). Increased pigmentation in *Galleria* is likely due to melanization of host tissues, and is indicative of an immune response to infectious agents [33]. Less pigmentation was observed in larvae that received smaller inoculums (10^4^ or 10^5^ spores) and was completely absent in PBS-injected controls. Thus, pigmentation was a specific response to fungal spores and not due to physical injury caused by the injection.

**Figure 1.**
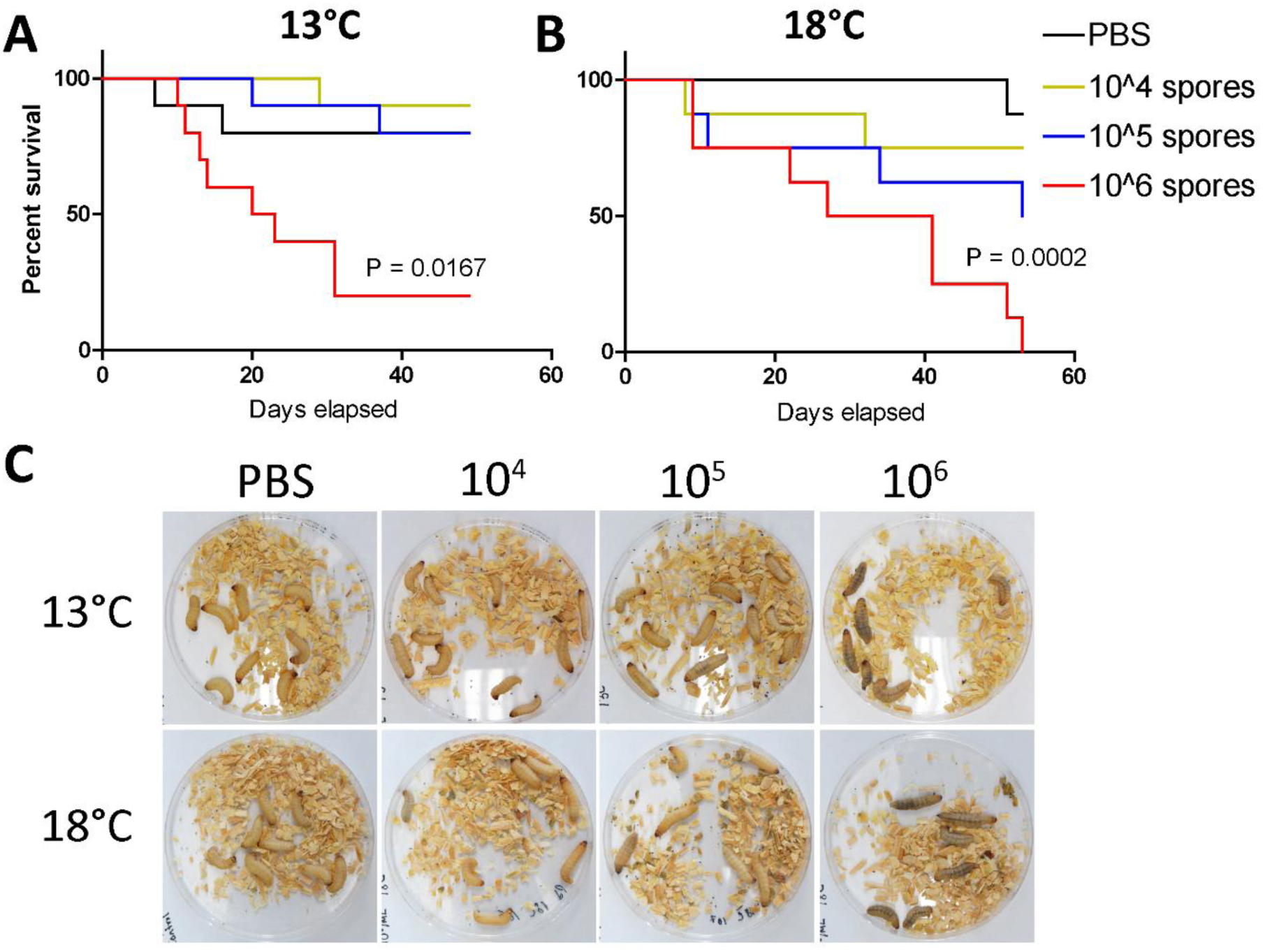
Inoculation of *G. mellonella* larvae with *P. destructans* spores leads to larval killing in an inoculum- and temperature-dependent manner. (A,B) 10 larvae each were injected with 10^4^, 10^5^ or 10^6^ spores or an equal volume of PBS (control) and kept at 13°C (A) or 18°C (B). Worms were monitored daily and deaths recorded. p-values represent Log-rank test in infections with 10^6^ spores v. PBS control. (C) Images of infected worms 16 days post-inoculation. Melanization can be seen in infected larvae and increases with inoculum size.

### Lethality requires infection by live spores

The killing of *G. mellonella* by *P. destructans* spores could be due to active proliferation of fungal cells in the larvae, or due to the host response to spores regardless of whether the spores were viable. To distinguish between these possibilities, heat-killed spores were prepared by treatment at 65**°**C for 30 minutes. Plating of heat-treated spores to YPD medium established that less than 0.1% of the spores were viable (Fig. 2, inset). Heat-killed and live spores were used to infect *G. mellonella* and larval death was monitored daily. While some mortality was observed in larvae infected with heat-killed spores, live *P. destructans* spores caused significantly greater killing of the larvae (P < 0.0001) (Fig. 2). This result indicates that the majority of larvae killed by *P. destructans* required infection with live spores. This was further supported by the recovery of viable *P. destructans* cells upon homogenization and plating of newly killed larvae onto YPD medium.

**Figure 2.**
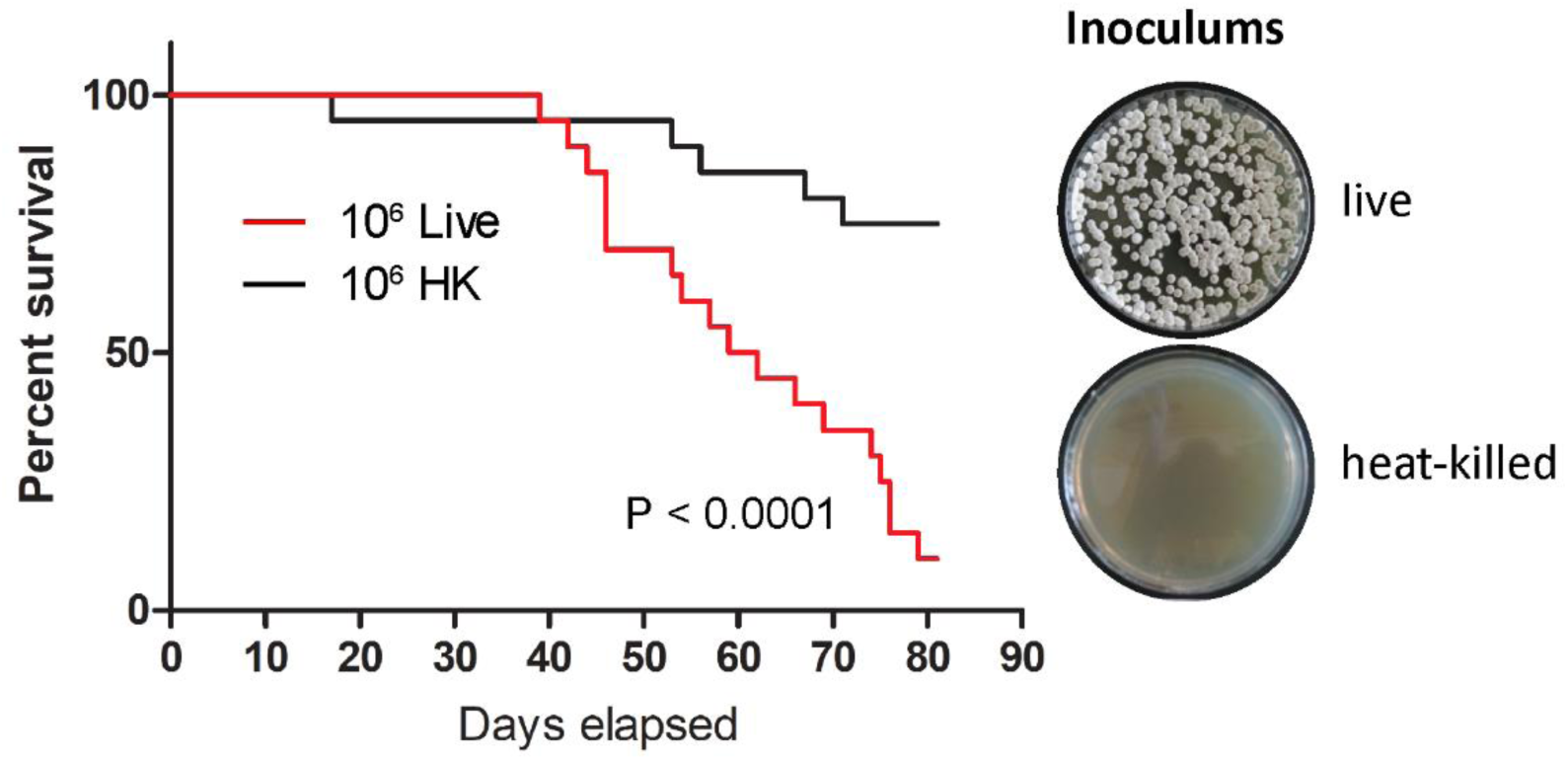
Effective killing of larvae requires live *P. destructans* spores. Larvae were injected with 10^6^ live spores or an equal number of heat-killed (HK) spores, kept at 13°C and monitored daily. 20 larvae were used per condition in two independent experiments, p-value represents result of Log-rank test on Live vs. HK. Right inset: representative YPD agar plates on which 1000 spores from live and heat-killed inoculums were plated to confirm efficacy of heat-killing.

### Virulence of *P. destructans* is increased by pre-germinating fungal spores

We examined whether spore germination could have an impact on the survival times of larvae. *P. destructans* spores were harvested and incubated at 13**°**C in liquid YPD medium for varying amounts of time to allow germination prior to infection of *G. mellonella*. Spores were cultured for 0, 6, 12 or 24 hours in YPD and microscopic examination revealed that germ tube formation was clearly visible in those grown for 12–24 hours (Fig. 3A). The extent of germination in each of the inoculums was calculated by counting spores with visible germ tubes versus non-germinated spores. We found that ~50% of spores incubated for 24 h in YPD had formed germ tubes and many cells had already formed hyphae (Fig. 3A and 3B). This contrasts with ~35% of spores having germinated after 12 h in YPD, while at 0 h or 6 h less than 10% of spores had visible germ tubes.

**Figure 3.**
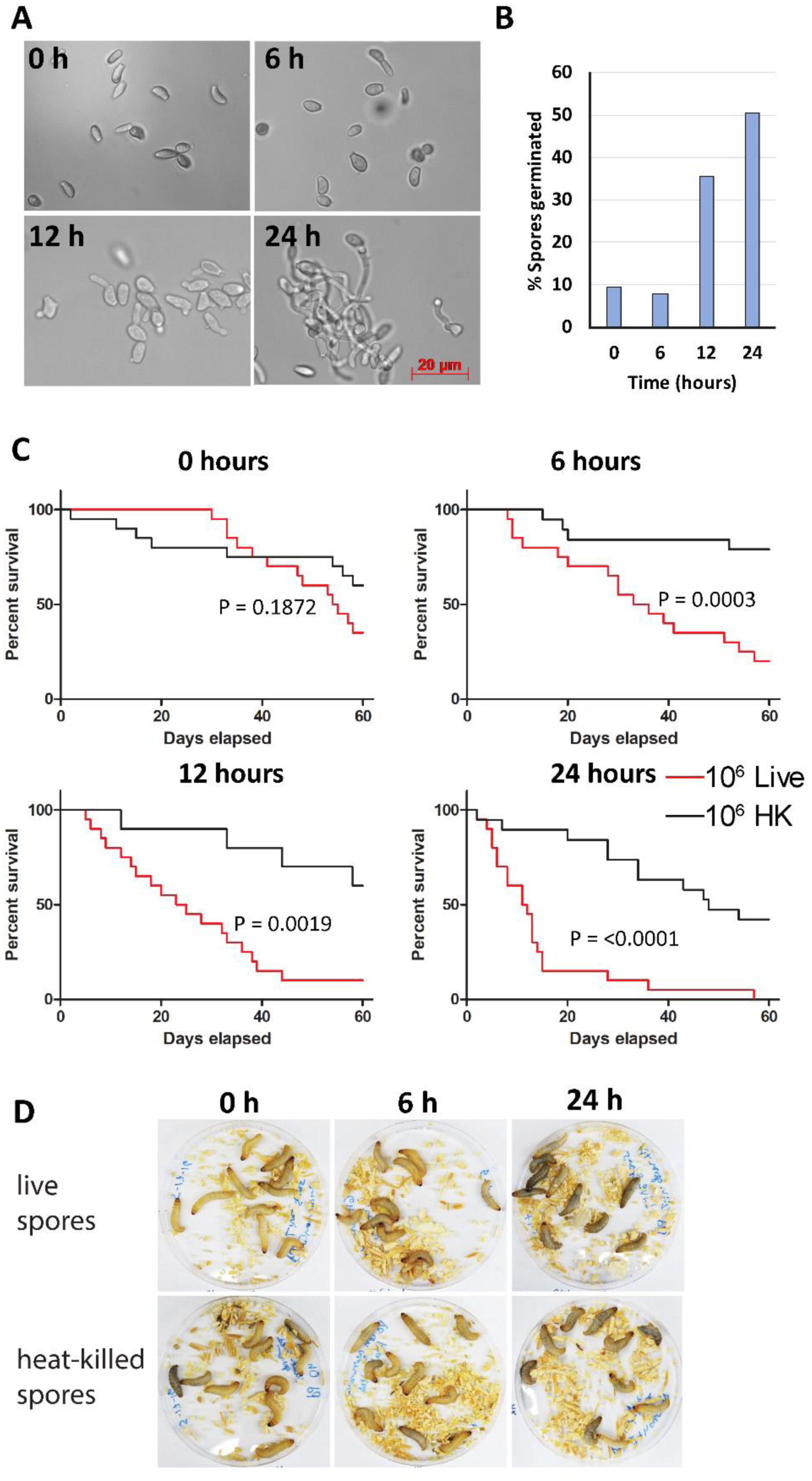
Pre-germinated *P. destructans* spores kill larvae more effectively than non-germinated spores. (A) Representative images of inoculums demonstrate formation of germ tubes by fungal spores after 12 and 24 h in YPD. (B) Quantification of germ tube formation at each time point. Data represents analysis of >= 400 spores from two independent experiments for each time-point. (C) Larvae were each injected with either 10^6^ live or heat-killed (HK) spores which had been allowed to pre-germinate for 0, 6, 12 or 24 h. Larval deaths were recorded daily. 20 larvae from two independent experiments were used per condition. p-values represent results of Log-rank test on live v. HK spores. (D) Images of worms infected with spores allowed to germinate for 0, 6, or 24 h (images taken 2 h post-infection) shows greatest melanization in larvae infected with spores pre-germinated for 24 h.

Germinated *P. destructans* spores were found to kill larvae much more rapidly than non-germinated spores. Live spores that had been pre-incubated for 24 hours at 13°C killed 50% of infected larvae within ~10 days, whereas larvae injected with spores germinated for 0, 6 or 12 h reached 50% mortality at approximately 45 days, 20 days or 15 days, respectively (Fig. 3C). Increased melanization was also observed in larvae infected with spores that were pre-germinated (both live and heat-killed), indicating that germinated spores may elicit a stronger immune response from the host (Fig. 3D). However, the increased host response did not directly impact mortality rates as germinated heat-killed spores still produced little death in the host.

### Antifungal drug screen with Phenotype Microarray (PM) plates

We next examined the feasibility of using *G. mellonella* larvae for evaluating antifungal agents against *P. destructans*. An *in vitro* screen was conducted to identify chemical inhibitors of *P. destructans* using the Phenotype Microarray (PM) system (Biolog) which consists of 96 well plates (PM21–25) coated with a panel of 120 chemical compounds at 4 different concentrations (for a full list of compounds see Supplementary Table 1)[29]. *P. destructans* spores suspended in YPD were used to inoculate each PM well and growth was monitored over the course of 11 days by measuring optical density (OD595). The resulting growth curves were analyzed using DuctApe software to generate an “activity index” representing relative fungal growth in the presence of a given compound (0 = no growth to 9 = maximal growth)[30]. Of the 120 compounds tested, approximately one third (42/120) were able to fully inhibit growth of *P. destructans* (activity index = 0) under at least one of the concentrations tested (Fig. 4A). The compounds that were able to fully inhibit growth are predicted to act on a wide range of cellular targets including the cell membrane, cell wall, protein synthesis and cellular respiration (Fig. 4B, Supp. Table 1). However, two prominent classes of inhibitory compounds included those targeting the cell membrane (p < 0.0001, Fisher’s exact test) and antipsychotics/efflux pump inhibitors (p = 0.0136), suggesting that these may represent aspects of *P. destructans* biology that are particularly sensitive to perturbation.

**Figure 4.**
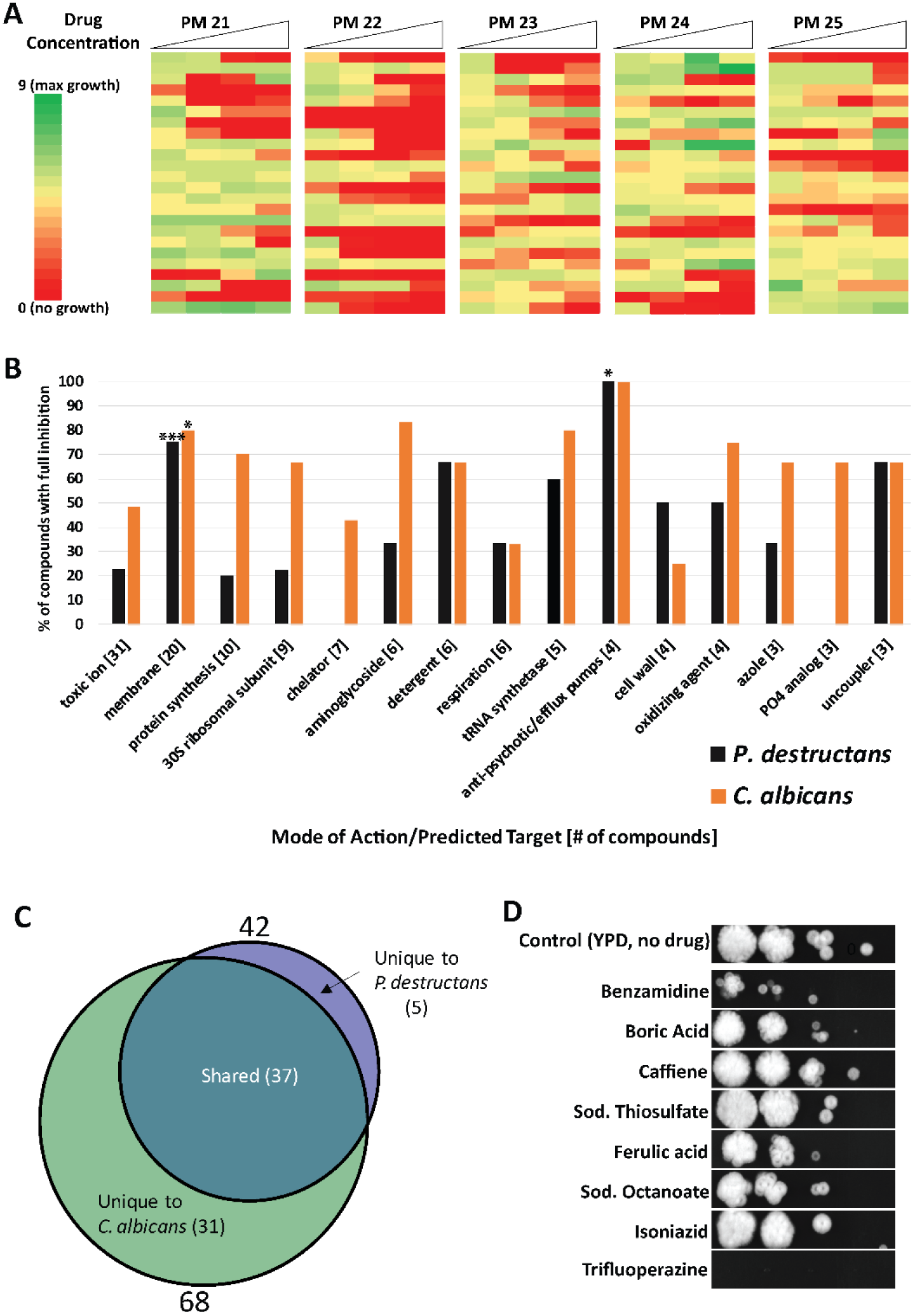
Identification and evaluation of anti-*P. destructans* compounds. (A) Heat map representing relative *P. destructans* growth on PM plates 21–25 which contain a total of 120 test compounds. (B) Bar graph indicating the % of PM compounds within each mode of action/target category that were able to fully inhibit growth of *P. destructans* (black) or *C. albicans* (orange) * P <= 0.05, *** P <= 0.001 (Fisher’s exact test). (C) Venn-diagram indicating compounds able to inhibit *P. destructans* and *C. albicans* growth. (D) Images of spores spotted onto YPD agar as 10-fold dilutions from left (10^3^) to right (10^0^) and allowed to grow for 1 week after an initial 24 h exposure to the indicated compound at 10 times the MIC concentration.

The sensitivity of *P. destructans* to chemical inhibitors was compared to that of the human fungal pathogen, *Candida albicans*, which was previously evaluated in PM plate assays (Fig. 4B, orange bars,[31]). *C. albicans* cells similarly showed a significant enrichment for inhibitory compounds targeting the cell membrane (p = 0.0260). Additionally, all four of the compounds annotated as efflux pump inhibitors within the PM panel (trifluoperazine, thioridazine, promethazine and chlorpromazine) were able to fully inhibit growth of both *P. destructans* and *C. albicans*. This indicates that the sensitivity of *P. destructans* to compounds targeting the cellular membrane and efflux pumps is not unique among fungal pathogens. This comparison also revealed that *P. destructans* was sensitive to a smaller number of PM compounds than *C. albicans* (42 and 68 compounds, respectively; Fig. 4C). Only 5 inhibitory compounds were unique to *P. destructans* and these were benzamidine, cadmium chloride, ceftriaxone, sodium azide and sodium thiosulfate. This may indicate that *P. destructans* is generally more resistant to chemical inhibitors than *C. albicans*, although the growth media differed between these two studies which limits direct comparisons between the two species.

### Evaluation of PM drug screen hits

A subset of the 42 inhibitory compounds identified from the PM screen were selected for further evaluation. Compounds known to possess high toxicity toward animal species were avoided. For each test compound, the minimum inhibitory concentration (MIC) was determined by testing a range of drug concentrations against *P. destructans* grown in YPD using the broth dilution method. The MIC of each compound was defined as the minimum concentration at which at least 80% of fungal growth was inhibited. The majority of compounds tested showed MICs within the millimolar range, however two compounds, trifluoperazine and sodium thiosulfate, had MICs in the micromolar range (130 µM or 53 µg/mL and 12.5 µM or 2 µg/mL, respectively) indicating they are relatively potent inhibitors of *P. destructans* growth (Table 1).

**Table 1.** Selected anti-*P. destructans* compounds from PM drug screen.

| **chemical** | **mode of action/targets** | **MIC value**<br>**(80% inhibition)** |
| --- | --- | --- |
| **Caffeine** | cyclic AMP phosphodiesterase inhibitor | **8 mM** |
| **Boric acid** | toxic anion | **25 mM** |
| **Benzamidine** | peptidase inhibitor, fungicide | **25 mM** |
| **Trifluoperazine** | membrane, phenothiazine, efflux pump inhibitor, anti-psychotic | **130 μM** |
| **Sodium Thiosulfate** | toxic anion, reducing agent | **12.5 μM** |
| **Ferulic acid** | antioxidant | **6 mM** |
| **Sodium Caprylate** | respiration, ionophore, H<sup>+</sup> | **50 mM** |
| **Isoniazid** | inhibitor of fatty acid synthesis | **11 mM** |

Each test compound was also evaluated to determine if it was fungistatic or fungicidal. To determine fungicidal activity, *P. destructans* spores were exposed to test compounds at a concentration 10-fold higher than the determined MIC for 24 h. Spores were then plated onto YPD to assess their viability (Fig. 4C). A compound was considered fungicidal if *P. destructans* was unable to grow on YPD plates after exposure. Of the compounds tested, only trifluoperazine showed fungicidal activity whereas all other compounds appeared fungistatic.

### Inhibitory compounds can block killing of *G. mellonella* larvae by *P. destructans*

Trifluoperazine and sodium thiosulfate, the most potent inhibitors of *P. destructans* identified *in vitro*, were tested for their ability to block killing of *G. mellonella* larvae. Two widely used antifungal drugs, amphotericin B and fluconazole [34], were also tested in these experiments, and we note that *P. destructans* has previously been shown to be susceptible to amphotericin B and dose-dependent susceptible to fluconazole [10]. For each assay, *P. destructans* spores were harvested, pre-germinated, resuspended in PBS containing each test compound and injected into *Galleria* larvae alongside controls consisting of spores in PBS alone. Deaths were recorded daily and viability curves constructed (Fig. 5 A-D). Treatment of spores with sodium thiosulfate or fluconazole failed to reduce larval killing compared to infections with untreated spores (Fig. 5A and 5B). However, both trifluoperazine and amphotericin B blocked larval killing by *P. destructans*, reducing the rate of mortality to that produced by heat-killed spores (p < 0.0001, log-rank test, Fig. 5C and 5D). In these experiments, untreated spores killed 50% of larvae after 20 days and 100% of larvae by 42 days, whereas >90% of larvae inoculated with trifluoperazine-treated spores and 65% of larvae inoculated with amphotericin B-treated spores were still viable 60 days post-infection. Amphotericin B, like trifluoperazine, is a fungicidal drug [35], whereas fluconazole is considered a fungistatic inhibitor [36]. These results suggest that fungicidal inhibitors were particularly effective in protecting *G. mellonella*.

**Figure 5.**
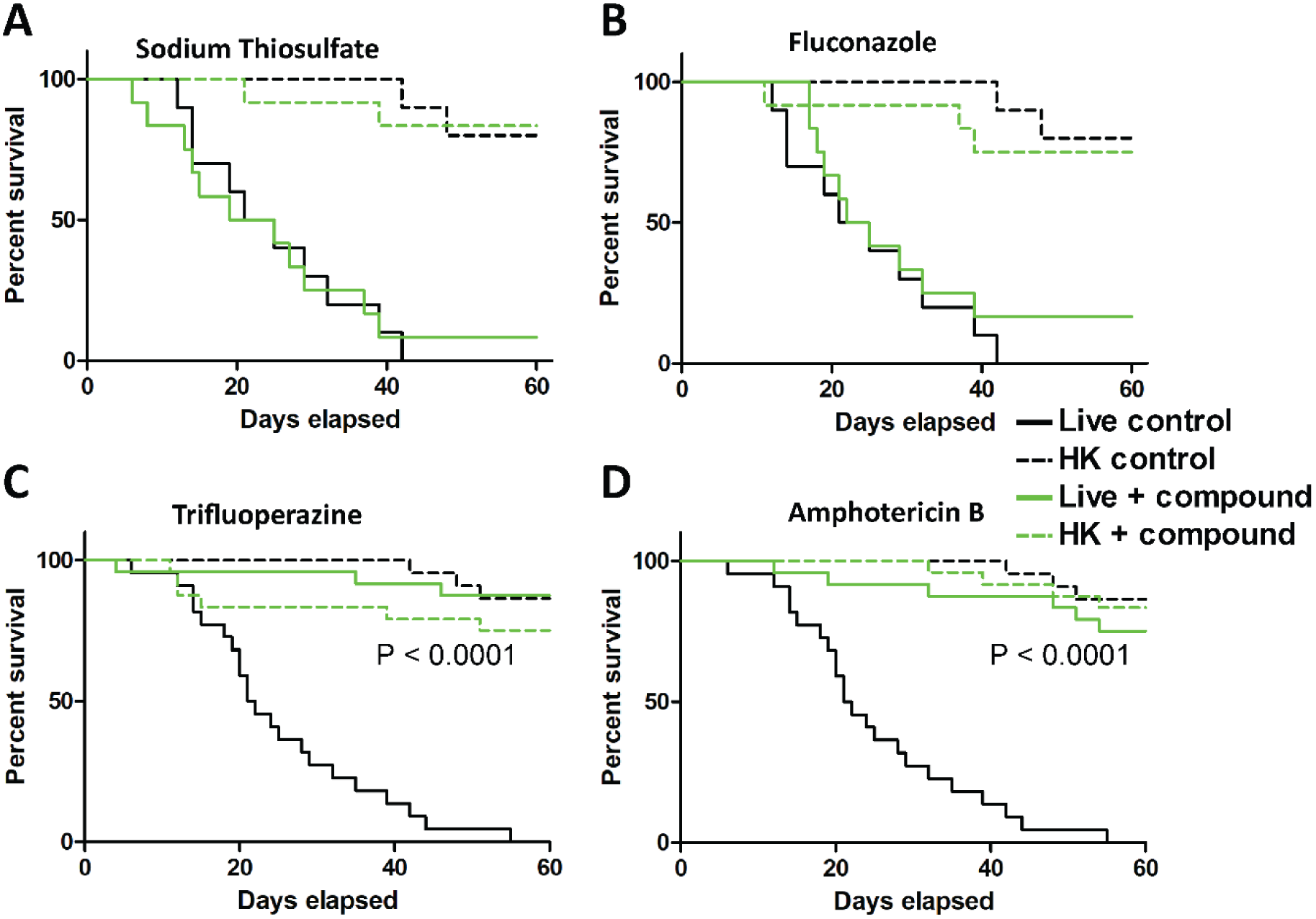
Evaluation of anti-*P. destructans* compounds using the *G. mellonella* infection model. Survival curves from *Galleria* larvae injected with spores (10^6^/worm) resuspended in PBS containing sodium thiosulfate (125 µM) (A), fluconazole (100 µg/ml) (B), trifluoperazine (1.3 mM) (C), or amphotericin B (200 µg/ml) (D). Each curve is plotted against control larvae injected with live or heat-killed (HK) spores germinated for 6 h and resuspended in PBS. p-values represent Log-rank test on live-treated vs. live-untreated spores.

Overall, our experiments demonstrate the utility of the *Galleria* model for the evaluation of *P. destructans* inhibitors and provide a proof-of-principle for screening of potential WNS treatments through the validation of trifluoperazine and amphotericin B as inhibitors of pathogenesis.

## Discussion

In this work, we establish the invertebrate *Galleria mellonella* as a suitable model for examining the virulence of *P. destructans*, the pathogen responsible for WNS. Our experiments demonstrate that *G. mellonella* larvae are susceptible to *P. destructans* at temperatures compatible with the growth of this fungus (13°C or 18°C). To our knowledge, this is the first evidence that *G. mellonella* can be used to study the virulence of cold-loving, psychrophilic species. Infection with *P. destructans* proceeds slowly, with larvae typically succumbing to infection over the course of 1–2 months. This time-frame is similar to infections of hibernating bats, where WNS gradually leads to progressive tissue damage and depletion of host reserves [8, 9].

*P. destructans* caused larval killing in an inoculum-dependent manner, with the most effective inoculum being 10^6^ spores per larva. These inoculums are similar to those described for other fungal pathogens in *G. mellonella* [20], although the length of the infection required for killing by *P. destructans* is considerably longer. Experiments using heat-killed *P. destructans* spores revealed that larval killing requires active fungal proliferation and is not simply a consequence of collateral damage from the host response to the inoculum. This is relevant as dead spores from some fungal species can kill larvae even in the absence of a live infection [37]. Together, these findings support the use of *G. mellonella* larvae as an accessible system for studying pathogenicity during infection by *P. destructans*.

We found that pre-germinating *P. destructans* spores had a considerable effect on virulence levels, as spores germinated for 24 hours prior to infection were able to kill larvae ~3–4 times faster than non-germinated spores. There may be several non-mutually exclusive factors contributing to this phenomenon. One possibility is that spore germination does not readily occur within the larvae and represents a rate-limiting step in the infection process. Pre-germinated spores may therefore exhibit an advantage as they have progressed past this rate-limiting step. Alternatively, germinated spores could be more resistant to phagocytosis or other immune defenses than non-germinated spores. In line with this, cell shape and/or size has been shown to determine interactions between some fungal species and immune cells, with elongated hyphal cells being more resistant to phagocytosis than smaller yeast cells [38]. *P. destructans* spores that have initiated hyphae formation may therefore present a greater challenge to clearance by larval hemocytes. Remodeling of the fungal cell wall during spore germination may also impact immune responses. *C. albicans* hyphal cells, but not yeast cells, can block phagosome maturation and induce macrophage lysis due to expression of hyphal-specific cell wall components [39, 40]. It is possible that a similar mechanism allows germinated *P. destructans* spores to resist the function of *G. mellonella* hemocytes, thereby accelerating infection. On the other hand, conidia from *Aspergillus fumigatus* possess hydrophobins [41] and melanins [42] that mask immunogenic components of the cell wall. Upon germination, *A. fumigatus* cells shed these components exposing surfaces that stimulate immune cell activation. The increased melanization of larvae injected with germinated spores suggests that *P. destructans* germ tubes may similarly stimulate a stronger immune response than non-germinated spores. However, given that germinated *P. destructans* spores are more virulent than non-germinated spores, this increased response is not protective and could even be detrimental to long-term survival in this model.

In addition to establishing conditions for infection of *G. mellonella* with *P. destructans*, we also used this model to test whether inhibitors of fungal growth could improve infection outcomes. A screen using Biolog PM plates identified 42 compounds that inhibited *P. destructans in vitro*, with effective inhibitors significantly enriched in compounds targeting the cell membrane and efflux pumps. A comparison with data from *C. albicans* [31] indicated that this species is similarly sensitive to these classes of inhibitors, although differences in growth conditions make it difficult to precisely compare inhibitor responses between fungal species. We note that Chaturvedi *et al*. previously performed a more extensive screen of 1,920 compounds of which 1.4% were highly effective at inhibiting *P. destructans* growth, including several azole drugs, a fungicide (phenylmercuric acetate), as well as several biocides [10]. The fraction of effective compounds identified was similar to that of high-throughput screens for compounds against other fungal species, whereas the high hit rate in our more limited screen (35%) is presumably due to Biolog compounds having been pre-selected for those with potential antifungal activity.

We subsequently focused our analysis on trifluoperazine and sodium thiosulfate from the Biolog screen, as both compounds showed inhibition of *P. destructans* in the micromolar range. Trifluoperazine is an antipsychotic drug that has been shown to have antimicrobial properties against a wide range of fungal and bacterial species [43, 44], whereas sodium thiosulfate has been used as a topical treatment for fungal infections [45]. We compared the action of these inhibitors with two well-established antifungal drugs, amphotericin B and fluconazole. Of these compounds, only trifluoperazine and amphotericin B effectively blocked larval killing by *P. destructans*. Both of these compounds are primarily fungicidal (our data and [35]), suggesting this mode of action may be most effective for this fungus. Curiously, trifluoperazine was also tested as part of the high-throughput screen performed by Chaturvedi *et al*. but was not found to inhibit *P. destructans* growth [10], which could be due to differences in the drug concentrations or culture conditions used. The use of fungicidal drugs would also be beneficial for treatment of WNS in nature, as delivery of a single application to hibernating bat populations would be considerably more practical than multiple applications. The *G. mellonella* model can therefore enable screening of anti-*P. destructans* compounds and may help accelerate the development of an effective WNS treatment.

Finally, we note that the *G. mellonella* model also has limitations that should be considered. First, WNS is a mammalian disease in which infection follows the invasion of external dermal tissues, whereas *Galleria* is an invertebrate species where fungal spores are directly introduced into the insect hemolymph. The larval model therefore lacks important features that occur during fungal colonization and invasion of dermal tissues. Second, *G. mellonella*, like all insects, lacks an adaptive immune system. However, this difference may not be critical for modeling *P. destructans* infections, as overall immune function is down-regulated in hibernating animals [46], and adaptive immune responses are particularly lacking in hibernating bats, whereas innate pro-inflammatory signaling is activated in response to *P. destructans* [26–28]. Neutrophils are also present in hibernating bats [27, 28, 47], although neutrophil recruitment and activation by *P. destructans* may primarily occur upon a return to euthermia [48]. A number key aspects of innate immunity are, in fact, shared between *G. mellonella* and mammals, including pathogen-associated molecular pattern (PAMP) receptors, anti-microbial peptides and phagocytic cells [20]. *Galleria* hemocytes share similarities with mammalian neutrophils and can phagocytose microbes, generate reactive oxygen species (ROS) and produce extracellular net-like structures for microbial killing [23–25]. The *G. mellonella* immune system, while lacking adaptive immunity, therefore shows several parallels to that of the natural host for *P. destructans*.

In conclusion, we demonstrate that *G. mellonella* represents a highly accessible model for the analysis of *P. destructans*, the primary cause of WNS. While there are limitations to this model its simplicity, ease of use, and affordability make it an attractive system for high-throughput screening of antifungal agents, as well as for the analysis of fungal mutants that may be defective in virulence. We suggest its inclusion will add to the growing list of important tools available for the study of this emerging mammalian pathogen.

## Acknowledgements

We would like to thank Iuliana Ene for assistance with PM data analysis, Corey Frazer and Iuliana Ene for comments on the paper, and Matthew Anderson and Matthew Hirakawa for help with larval injections.

## Funding

Funding for this project was provided by a National Science Foundation grant (NSF-1456787) to RJB. The funders had no role in study design, data collection and analysis, decision to publish, or preparation of the manuscript.

